# T cell subset-selective *IL2RA* enhancers shape autoimmune diabetes risk

**DOI:** 10.1101/2020.07.22.216564

**Authors:** Dimitre R. Simeonov, Harikesh S. Wong, Jessica T. Cortez, Arabella Young, Zhongmei Li, Vinh Nguyen, Kyemyung Park, Jennifer Umhoefer, Alyssa C. Indart, Jonathan M. Woo, Mark S. Anderson, Ronald N. Germain, Alexander Marson

## Abstract

The majority of genetic variants associated with complex human autoimmune diseases reside in enhancers^1–3^, non-coding regulatory elements that control gene expression. In contrast with variants that directly alter protein-coding sequences, enhancer variants are predicted to tune gene expression modestly and function in specific cellular contexts^4^, suggesting that small alterations in the functions of key immune cell populations are sufficient to shape disease risk. Here we tested this concept by experimentally perturbing distinct enhancers governing the high affinity IL-2 receptor alpha chain (IL2RA; also known as CD25). IL2RA is an immune regulator that promotes the pro- and anti-inflammatory functions of conventional T cells (Tconvs) and regulatory T cells (Tregs), respectively, and non-coding genetic variants in *IL2RA* have been linked to multiple autoimmune disorders^4^. We previously tiled across the *IL2RA* locus using CRISPR-activation and identified a stimulation-responsive element (CaRE4) with an enhancer that modestly affects the kinetics of IL2RA expression in Tconvs^5^. This enhancer is conserved across species and harbors a common human SNP associated with protection from Type 1 Diabetes (T1D)^5,6^. We now identified an additional conserved enhancer, termed CaRE3 enhancer, which modestly affected steady state IL2RA expression in regulatory T cells (Tregs). Despite their seemingly subtle impact on gene expression, the CaRE3 and CaRE4 enhancers had pronounced yet divergent effects on the incidence of diabetes in autoimmune prone animals. Deletion of the conserved CaRE4 enhancer completely protected against autoimmune diabetes even in animals treated with an immunostimulating anti-PD1 checkpoint inhibitor, whereas deletion of the CaRE3 enhancer accelerated spontaneous disease progression. Quantitative multiplexed imaging of the pancreatic lymph nodes (panLNs) revealed that each enhancer deletion preferentially affected the protein expression levels of IL2RA in activated Tconvs or Tregs, reciprocally tuning local competition for IL-2 input signals. In animals lacking the CaRE4 enhancer, skewed IL-2 signaling favored Tregs, increasing their local density around activated Tconvs to strongly suppress emergence of autoimmune effectors. By contrast, in animals lacking the CaRE3 enhancer, IL-2 signals were skewed towards activated Tconvs, promoting their escape from Treg control. Collectively, this work illustrates how subtle changes in gene regulation due to non-coding variation can significantly alter disease progression and how distinct enhancers controlling the same gene can have opposing effects on disease outcomes through cell type-selective activity.

---

Given the central role in immune homeostasis played by IL2RA expression and IL-2 signaling in distinct subpopulations of CD4^+^ T cells^7^, we sought to understand how distinct *IL2RA* enhancers regulate expression of IL2RA in CD4^+^ Tconvs and in Tregs to influence the risk of autoimmunity. Previously, we identified a conserved intronic *IL2RA* enhancer, termed the CaRE4 enhancer, which tunes the kinetics of IL2RA induction in Tconvs following T cell receptor (TCR)-mediated stimulation but had only subtle effects on IL2RA levels in FOXP3^+^ Tregs at steady-state^5,8^ (Fig. 1a). Given their unique transcriptional circuitry and functional requirement for IL-2 signals, we hypothesized that Tregs use enhancers distinct from Tconv to maintain high IL2RA expression at steady-state.

**Figure 1.**
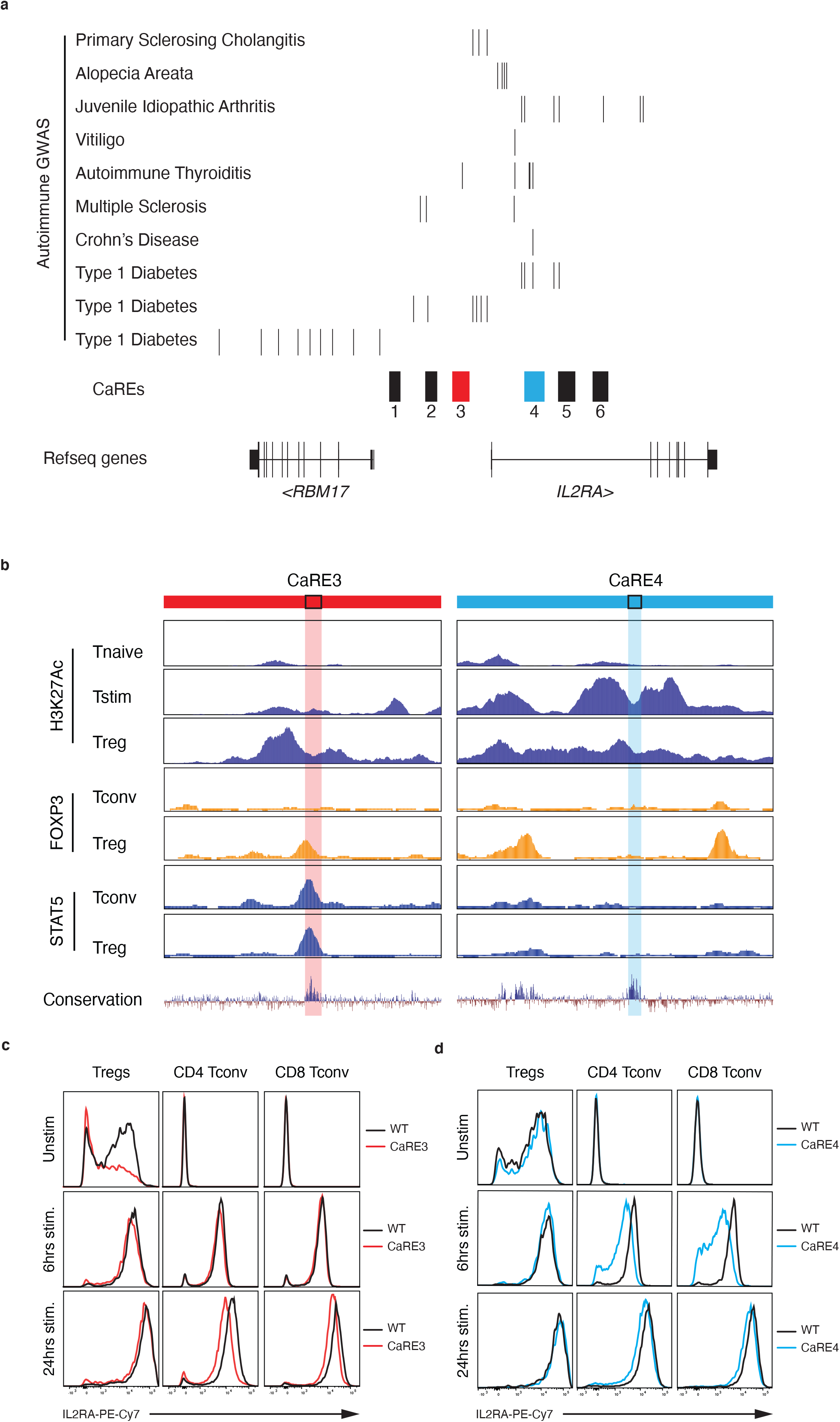
Mapping the functional immune contexts of *IL2RA* CaRE3 and CaRE4 enhancers. **a**, The *IL2RA* locus showing fine-mapped candidate autoimmunity variants (T1D^6^, Crohn’s Disease^15^, and others^4^) and CRISPRa responsive elements (CaREs)^5^, including CaRE3 and CaRE4. **b**, *IL2RA* CaRE3 (hg19 chr10:6109198-6113209) and CaRE4 (hg19 chr10:6092205-6096760) overlap with ChIP-seq for H3K27Ac (Epigenome Roadmap Project) and the transcription factors FOXP3 and STAT5 in subsets of primary human T cells^9,23^. The lighter color vertical bars represent the conserved enhancers investigated in this study. STAT5 and FOXP3 ChIP performed on *in vitro* expanded Tconv and Treg sorted as CD4+CD45RA+IL2RA− and CD4+CD45RA+IL2RAhigh, respectively. H3K27Ac signal y-axis range 2-120 for CaRE3 and 2-300 for CaRE4. STAT5 and FOXP3 ChIP signal y-axis range 1-40 for CaRE3 and CaRE4. **c, d,** IL2RA surface expression on T cell subsets stimulated in splenocyte cultures from NOD CaRE3 (n=1) or CaRE4 EDEL (n=1) mice versus WT littermate control. Data from NOD mice with homozygous CaRE3 deletion in red, homozygous CaRE4 deletion in blue, and WT littermate controls in black.

We searched for candidate enhancer elements that appeared preferentially active in Tregs by analyzing histone 3 lysine 27 acetylation (H3K27Ac) data^9^ from subsets of primary human T cells. This approach identified a conserved ~200bp sequence upstream of the *IL2RA* promoter, where associated histones (H3) were highly acetylated (K27) in Tregs but not in Tconvs (Fig. 1b). This site falls within a CRISPRa responsive element (CaRE) that affected IL2RA expression in our previous tiling screen and is referred to as the CaRE3 enhancer^5^ (Fig. 1a). H3K27Ac at this region was unaffected by TCR-mediated stimulation in Tconvs, unlike the CaRE4 enhancer (Fig. 1b). To further assess cell-type specific effects, we compared transcription factor binding between the CaRE3 and CaRE4 enhancers in Tregs and Tconvs. This analysis revealed that FOXP3 and STAT5 selectively bound to the CaRE3 enhancer in human Tregs (Fig. 1b). Both of these transcription factors have been reported to directly promote *IL2RA* expression in Tregs ^10−12^. By contrast, motif analysis suggested that a MEF2 binding site exists in the CaRE4 enhancer. Interestingly, this DNA sequence motif is disrupted by the common human SNP associated with protection from T1D (Extended Data Fig. 1)^6^. These results suggested that the CaRE3 and CaRE4 enhancers can be occupied by distinct TFs and may preferentially regulate *IL2RA* expression in Tregs and activated Tconvs, respectively.

To determine the *in vivo* function of these two *IL2RA* enhancers we generated CaRE3 enhancer deletion (EDEL) mice in both C57BL/6 and non-obese diabetic (NOD) backgrounds and compared them to previously developed CaRE4 EDEL mice^5,8^. The founder lines were backcrossed at least one generation before being used for experiments; immunophenotyping was generally consistent across founders and genetic backgrounds. We first characterized effects of the CaRE3 EDEL on T cell development in C57BL/6 mice. Within the thymus, we found no significant changes in the abundance of the major thymocyte populations (double negative (DN), double positive, CD4 and CD8 single positive) as compared to WT animals although IL2RA expression on DN thymocytes was impaired (Extended Data Fig. 2 and 3). Next, we examined Treg development in CaRE3 EDEL animals. In the thymus, we found reduced numbers of mature Tregs (FOXP3^+^IL2RA^+^) along with changes in the abundance of Treg progenitor populations (Extended Data Fig. 2 and 3). Consistent with these findings, we observed impaired, but not completely ablated, IL2RA expression on many CaRE3 EDEL Tregs in the thymus and in all other lymphoid tissues that were immunophenotyped (Extended Data Fig. 2 and 3). The presence of a CaRE3 binding site for STAT5, a TF downstream of IL-2 signaling, suggests this element could help to maintain Treg IL2RA levels in response to IL-2 produced by rare Tconvs at steady state^13–17^. In summary, our data suggests that the CaRE3 enhancer is required to maintain high steady-state expression of IL2RA in the Treg lineage.

We next tested how the CaRE4 and CaRE3 enhancers influence IL2RA levels in response to strong T cell activation. For this purpose, splenocytes from WT, CaRE3 EDEL, and CaRE4 EDEL NOD mice were incubated with soluble anti-CD3/CD28 antibodies, and IL2RA surface expression was measured by flow cytometry at multiple time points after stimulation. Consistent with our previous findings^8,13^, CaRE4 EDEL Tconv cells had impaired IL2RA levels early after activation (6 hours), but eventually approached WT cells at later time points (24 hours) (Fig. 1c, Extended Data Fig. 3). This kinetic delay in IL2RA induction was not observed in activated CaRE3 EDEL Tconv cells, although we noted reduced IL2RA on these cells at later time points. These findings demonstrated that the CaRE3 enhancer plays a relatively minor role in the early induction of IL2RA on Tconv cells compared to the CaRE4 enhancer. We next compared IL2RA levels on resting and activated Tregs in the splenocyte cultures. In the absence or presence of stimulation, both WT and CaRE4 EDEL Tregs expressed comparable levels of IL2RA. By contrast, unstimulated CaRE3 EDEL Tregs showed reduced IL2RA surface expression versus WT counterparts, similar to our *in vivo* findings (Fig. 1c, Extended Data Fig. 2 and 3). This reduction in IL2RA surface expression was restored to WT levels following stimulation, both at early and late timepoints. Given the known control of IL2RA expression by the strength of TCR signaling and IL-2-mediated positive feedback, both of these signals could contribute to this effect^14^. Thus, despite starting with impaired IL2RA levels, CaRE3 EDEL Treg cells retained stimulation-responsive IL2RA regulation, at least under conditions of strong T cell activation. Our data suggest that the CaRE4 enhancer is part of a stimulation-response program controlling the rate of IL2RA induction in Tconv. In these *in vitro* experiments, the CaRE3 enhancer was required for maintenance of high steady-state levels of IL2RA expression on Tregs.

Multiple autoimmune diseases have been linked to variants in non-coding sequences at the *IL2RA* locus. Notably, the non-coding SNP rs61839660, which resides within the CaRE4 enhancer, has been convincingly shown to influence autoimmunity risk, including promoting risk of Crohn’s Disease^15^. The variant is associated with protection against T1D^6^, although several other candidate SNPs are in linkage disequilibrium. We previously showed that the SNP impairs enhancer function and delays IL2RA expression in activated Tconvs^5^ (Fig. 1c and Extended Data Fig. 3). To understand how enhancer perturbations shape disease risk *in vivo,* we studied CaRE3 and CaRE4 deletion in NOD mice, which develop autoimmune diabetes with age and serve as a model for human T1D. Diabetes was assessed by monitoring blood glucose levels weekly. Whereas most WT NOD animals developed disease during the 30 week study, remarkably, all homozygous CaRE4 EDEL animals were protected completely from diabetes (Fig. 2a). This experimental result supports the putative protective role of this common human variant in T1D (Extended Data Fig. 1). Whereas the human single nucleotide variant and linked variants have a modest protective effect on T1D risk in population studies^6^, we discovered here that ablation of the whole conserved non-coding enhancer had a profound effect and completely prevented diabetes in the NOD mice. Consistent with observations that heterozygous human variants can modulate autoimmune disease risk^6,16^, we also observed reduced diabetes incidence in heterozygous animals that had the CaRE4 enhancer deleted on a single allele (Fig. 2a). Immunohistochemistry of pancreatic specimens revealed that homozygous CaRE4 EDEL mice had little to no immune infiltration in the islets, while heterozygous CaRE4 EDEL mice had intermediate infiltration compared to WT mice at 16 weeks of age (Extended Data Fig. 4). These perturbation studies of a non-coding element implicated in T1D risk by human genetic studies link the expression kinetics of *Il2ra* to autoimmune diabetes pathology.

**Figure 2.**
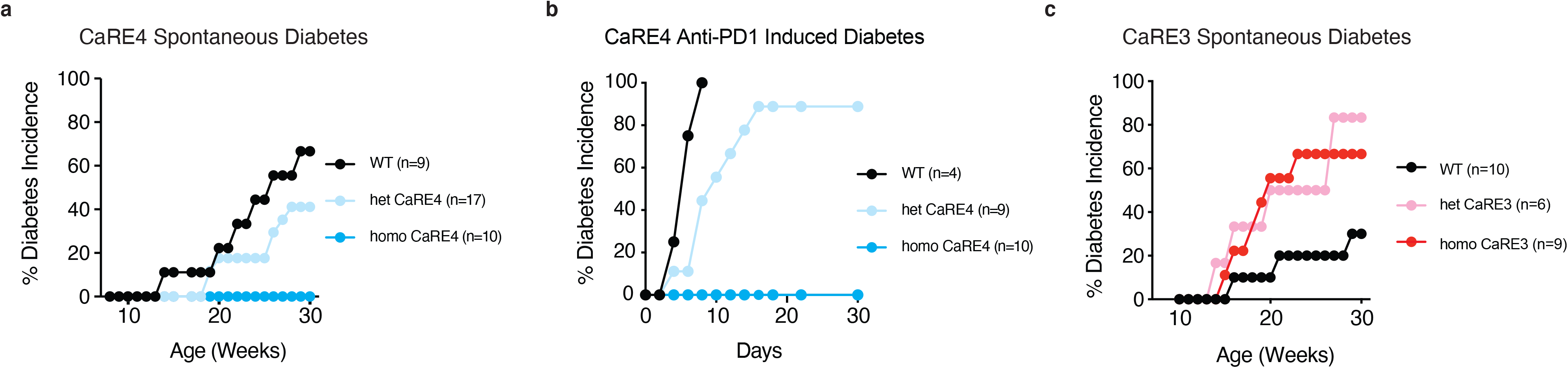
*Il2ra* CaRE3 and CaRE4 enhancers have divergent effects on autoimmune diabetes outcomes. **a**, Spontaneous diabetes incidence in NOD WT (n=9), heterozygous (n=17), and homozygous (n=10) CaRE4 EDEL female mice. **b**, Anti-PD1 induced diabetes incidence in NOD WT (n=4), heterozygous (n=9), and homozygous (n=10) CaRE4 EDEL mice. **c,** Spontaneous diabetes incidence in NOD WT (n=10), heterozygous (n=6), and homozygous (n=9) CaRE3 EDEL mice. Only female NOD mice were used for these studies.

We next tested if the CaRE4 element deletion could protect not only from spontaneous autoimmune diabetes, but also from induced diabetes due to acute challenge to immune tolerance by immunomodulatory therapies. For instance, blockade of the PD1-PDL1 checkpoint pathway is now widely used for cancer immunotherapies but induces autoimmune diabetes in a subset of patients^16,17^. Consistent with the toxicity observed in humans, PD1 blocking antibodies also cause acute onset diabetes in NOD mice^17^. We tested if a cohort of CaRE4 EDEL mice would be protected from diabetes after PD1 checkpoint blockade. Following treatment with PD1 blocking antibody none of the homozygous CaRE4 EDEL mice developed diabetes for the 30-day duration of the study, whereas all WT mice developed diabetes by day 10 of the study (Fig. 2b). Heterozygous deletion of the CaRE4 enhancer conferred intermediate protection with delayed onset of anti-PD1-induced diabetes (Fig. 2b). Histology from CaRE4 EDEL mice at the end of the study showed little to no islet immune infiltration even with anti-PD1 treatment (Extended Data Fig. 4e). Thus, deletion of the CaRE4 enhancer prevented the loss of immune tolerance caused by PD1 checkpoint blockade. These findings suggest that genetic variants mapping to cell-type specific IL2RA enhancers may be predictive of autoimmune side effects following cancer immunotherapy. Moreover, these findings emphasize how subtle changes in the expression kinetics of IL2ra can have pronounced effects on the progression of both spontaneous and induced forms of autoimmune diabetes.

In contrast to the protective effects we observed with CaRE4 deletion, deletion of the CaRE3 enhancer element exacerbated autoimmune diabetes in NOD mice (Fig. 2c). Homozygous or heterozygous deletion of the CaRE3 enhancer increased the progression to diabetes compared to WT littermate control mice, although we note an overall lower incidence of disease in this cohort of WT animals (Fig. 2c). These results demonstrated that perturbing the CaRE3 enhancer, even at a single *Il2ra* allele, was sufficient to accelerate the breakdown in immunological homeostasis.

We next investigated the mechanism by which the CaRE3 and CaRE4 enhancers in the same gene locus have divergent effects on autoimmune disease risk. We previously demonstrated that within LNs of healthy hosts, rare CD4^+^ T cells were activated by self-antigens (self-activated CD4^+^ Tconvs) and produced IL-2, which upon being sensed by neighbouring Tregs, initiated a local feedback process that prevented the emergence of tissue-damaging effector cells. Specifically, early IL-2 sensing by Tregs enhanced their local density and immunosuppressive capacity around self-activated T cells, thus constraining incipient autoimmune responses. Perturbing this local feedback process enabled self-activated T cells to respond to IL-2 and escape the control of Tregs, leading to a breakdown in immunological homeostasis^18,19^. Based on these prior observations, we hypothesized that disrupting the CaRE3 or CaRE4 enhancers would skew IL-2 signals in favor of self-activated CD4^+^ Tconvs or Tregs, respectively, and account at least in part for the altered disease risk of the EDEL NOD animals. This possibility was tested initially by comparing pSTAT5 expression between Tconvs and Tregs in the panLNs of WT or EDEL NOD animals. Within the panLN paracortex of WT NOD animals, the percentage of pSTAT5^+^ Tregs was much greater than that of pSTAT5^+^ CD4^+^ Tconv cells, as expected (Extended Data Fig. 5a). However, deletion of the CaRE3 enhancer resulted in a ~2-fold decrease in the percentage of pSTAT5^+^ Tregs and a corresponding ~3-fold increase in the percentage of pSTAT5^+^ CD4^+^ Tconvs (Fig. 3a, Extended Data Fig. 5a). In almost all instances, pSTAT5^+^ CD4^+^ Tconvs were highly enriched in PD1, which serves as a reliable readout of self-activation due to its rapid upregulation following TCR-mediated stimulation and high signal-to-noise ratio *in situ*^19^ (Fig. 3b). In contrast, deletion of the CaRE4 element strongly biased IL-2 signaling in favor of Tregs, resulting in a very small number of pSTAT5^+^ CD4^+^ Tconvs (Fig. 3a, Extended Data Fig. 5a). Thus, ablation of the *Il2ra* CaRE3 and CaRE4 enhancers respectively skewed the local competition for IL-2 in favor of rare self-activated CD4^+^ Tconvs or Tregs, thereby altering progression of autoimmune responses.

**Figure 3.**
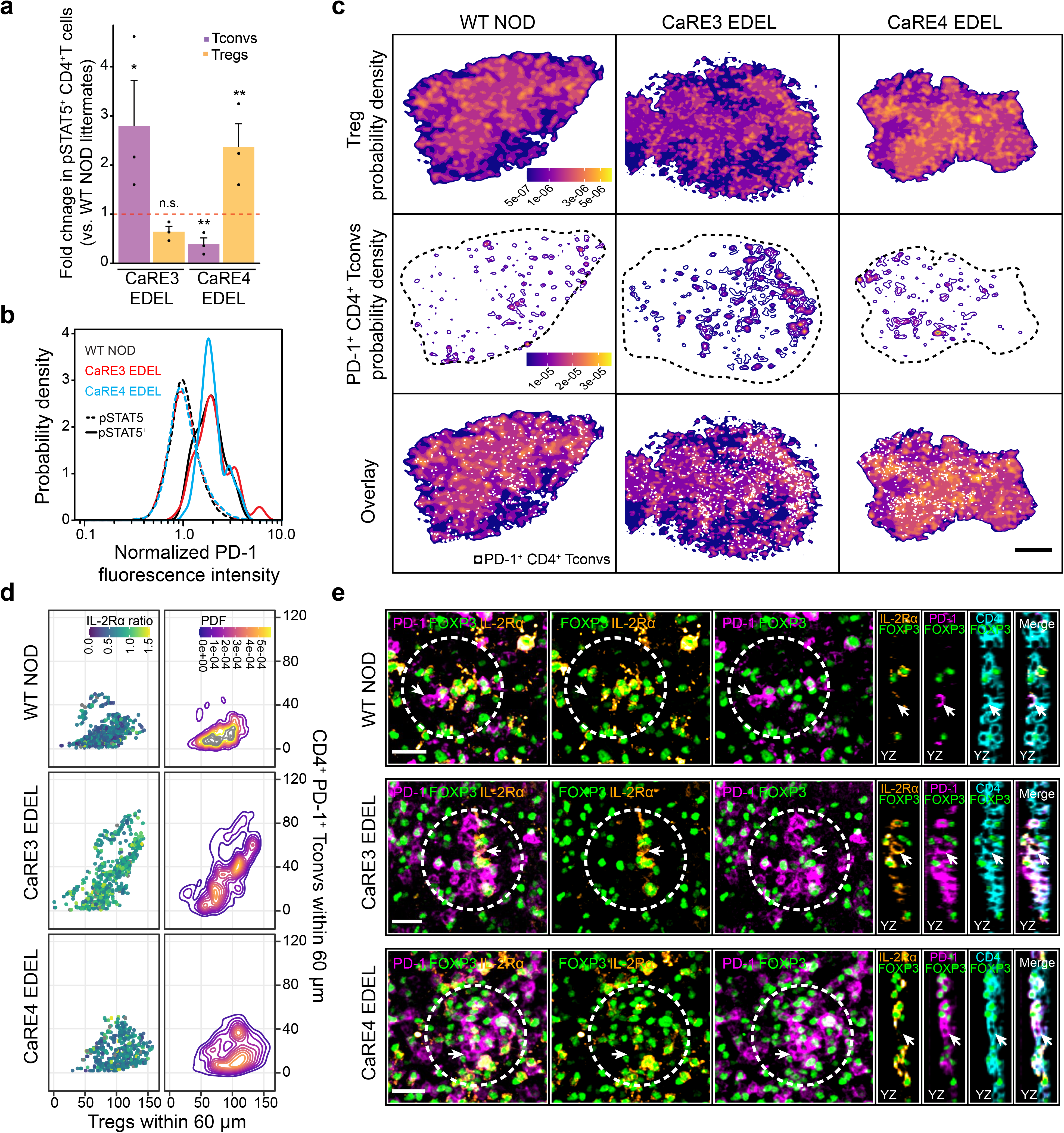
*Il2ra* enhancer deletions reciprocally tune the local competition for IL-2 between self-activated CD4^+^ Tconvs and Tregs. **a,** *In situ* quantification of pSTAT5+CD4+ Tconvs and pSTAT5+ Tregs within the panLN paracortex of 6-week-old female mice. pLNs were compared between CaRE3 and CaRE4 EDEL NOD (n=3) and WT NOD (n=3) littermate controls. Data are presented as mean +/− s.d. and are from two independent experiments. **b,** Representative probability density functions (PDFs) of PD1 expression in pSTAT5- or pSTAT5+CD4+ Tconvs in the panLNs of WT or EDEL NOD mice. PDFs were determined by kernel density estimation. **c**, Representative PDFs of Treg (top row) or PD1^+^CD4^+^ Tconv (middle row) XY spatial coordinates in the panLNs of WT NOD or EDEL NOD mice (see Methods). White dots represent the positions of individual PD1+CD4+ T cell within the paracortex (bottom row). Scale bar = 200 μm. **d,** *In situ* multiparameter analysis of the number of Tregs (x-axis) and CD4^+^ PD1^+^ Tconvs (y-axis) surrounding individual CD4+PD1+ Tconvs (individual dots). The IL2RA ratio was calculated by dividing the median IL2RA fluorescence intensity of each PD1+CD4+ Tconv by the median IL2RA fluorescence intensity of the surrounding Tregs within 60 μm. **e,** Representative confocal micrographs illustrating changes in the local density (dashed white circles) of Tregs and CD4+ PD1+ Tconvs as well as changes in the IL2RA expression ratio between WT NOD and EDEL NOD mice. YZ optical slices highlight CD4, PD1, and IL2RA expression on the cell of interest (white arrow). Scale bars = 25 μm.

Skewing of IL2 signals coincided with pronounced changes to the local densities of Tregs or self-activated CD4^+^ Tconvs throughout the panLN paracortex, consistent with the documented role of IL-2 in promoting T cell proliferation and survival (Fig. 3c)^7,20^. Indeed, in CaRE4 EDEL mice, we observed more Tregs surrounding individual PD1^+^ CD4^+^ Tconvs compared to WT or CaRE3 EDEL mice. This observation correlated with a reduced IL2RA expression ratio between individual PD1^+^ CD4^+^ T convs and proximal Tregs in CaRE4 EDEL mice (Fig. 3d, e and Extended Data Fig. 5b-d). Conversely, in CaRE3 EDEL mice, we observed dense clusters of PD1^+^ CD4^+^ Tconvs, suggesting that some self-activated Tconvs were escaping the control of Tregs and undergoing significant bursts of proliferation. While Tregs also exhibited elevated densities in several of these clusters, they did not appear to scale sufficiently to successfully constrain all of the self-activated CD4^+^ T cells (Fig. 3d, e). Consistent with this notion, the IL2RA expression ratio between individual PD1^+^ CD4^+^ T convs and proximal Tregs was greatly elevated when compared to WT NOD or CaRE4 EDEL mice (Fig. 3c and Extended Data Fig. 5). These results demonstrated that germline perturbations in *IL2ra* enhancers had cell type selective effects on gene expression and significantly imbalanced the local competition for IL-2 between self-activated CD4^+^ Tconvs and Tregs. Overall, our findings illustrate how critical non-coding *cis-*regulatory sequences in the *IL2RA* locus can empower either self-activated CD4^+^ Tconvs or Tregs and alter the risk of autoimmunity *in vivo*.

## Discussion

Here we show that distinct enhancers of the same gene can function in cell type selective contexts, thereby promoting divergent immunological consequences. In healthy hosts, this form of tight, context-selective gene regulation is likely critical for ensuring host protection while simultaneously limiting autoimmune tissue damage. Our data suggest that genetic variants can disrupt this delicate balance and alter the risk of autoimmune disease by perturbing enhancers with cell-type selective functions (Fig. 4). Even minor regulatory effects on gene expression appear to have strong non-linear effects on autoimmunity due to escape and expansion of rare self-activated Tconvs. These findings are consistent with the fact that non-coding sequences in the *IL2RA* locus have been linked to at least 8 different autoimmune disorders^4^. In addition, our *in vivo* data suggests that genetic variation in the *IL2RA* locus may help to predict risk of autoimmune disorders in response to anti-PD1 therapy or other forms of cancer immunotherapy. This hypothesis warrants further clinical study in humans undergoing checkpoint blockade treatments. Based on the data in this report, *IL2RA* enhancer variants may bias the competition for IL-2 between cell types by preferentially affecting either Tconvs or Tregs, leading to inappropriate immune responses^21,22^. Our current studies revealed effects of skewed IL-2 competition on peripheral tolerance, but effects on central tolerance may exist as well. Moreover, effects on additional cellular contexts with disease-specific relevance could contribute to the divergent associations of the rs61839660 SNP with Crohn’s disease risk and T1D protection. Looking forward, manipulating IL-2 competition by targeting the activated Tconv-selective CaRE4 enhancer or Treg-selective CaRE3 enhancer could open up new avenues for immunotherapies. Our current studies of the *in vivo* immunological effects of preferential Treg and Tconv enhancers will serve as a foundation for the rational engineering of non-coding circuits that promise to make the next generation of cellular immunotherapies safer and more effective. Overall, our results begin to define an enhancer code that shapes cell types, cellular functions, and biological outcomes.

**Figure 4.**
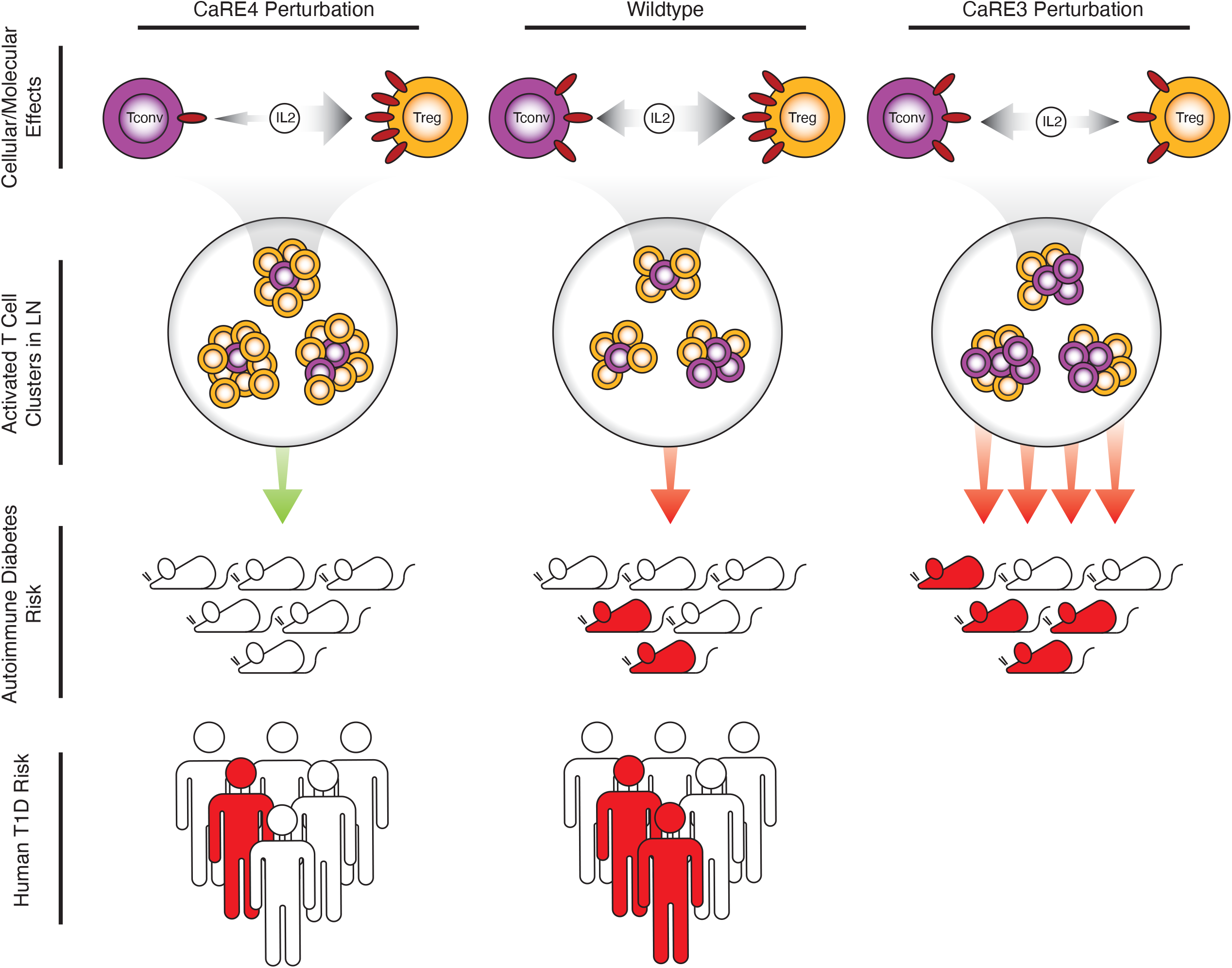
Model of *Il2ra* enhancer code and autoimmune diabetes risk. *Il2ra* CaRE3 and CaRE4 enhancers tune IL2RA levels in a cell-type selective manner on Tconvs or Tregs, respectively, skewing competition for IL-2. In the pancreatic lymph node, rare autoreactive Tconv cells compete with immunosuppressive Tregs for the consumption of the survival factor IL-2, which determines whether activated Tconv cells accumulate. Perturbation of the CaRE3 enhancer reduces IL2RA on Tregs, increases IL-2 signals in Tconv cells and promotes the formation of activated T cell clusters that are relatively deficient in immunosuppressive Tregs. In contrast, perturbation of CaRE4 impairs induction of IL2RA on Tconv cells, increases IL-2 signals in Tregs and promotes the formation of T cell clusters dominated by Tregs. Whereas CaRE3 deletion increased incidence of autoimmune diabetes, CaRE4 deletion completely protected against diabetes. Consistent with these findings, the conserved CaRE4 enhancer harbors a common human SNP associated with protection against T1D.

## Acknowledgements

We thank all members of the Marson and Germain labs. We thank Jeffrey A. Bluestone for his continued mentorship and guidance throughout the course of this work. D.R.S. was supported by the Jeffrey G. Klein Family Fellowship in Diabetes. J.T.C. is supported by the National Science Foundation Graduate Research Fellowship Program. This was work supported funds from NIH/NIDDK (DP3DK111914-01), the Chan Zuckerberg Biohub, the Northern California JDRF Center of Excellence and was supported in part by the Intramural Research Program of the National Institute of Allergy and Infectious Diseases, NIH. The Marson lab has received gifts from J. Aronov, G. Hoskin, K. Jordan, B. Bakar, the Caufield family and funds from the Innovative Genomics Institute (IGI), and the Parker Institute for Cancer Immunotherapy (PICI). A.M. holds a Career Award for Medical Scientists from the Burroughs Wellcome Fund, is an investigator at the Chan–Zuckerberg Biohub and is a recipient of a The Cancer Research Institute (CRI) Lloyd J. Old STAR grant. This work used the UCSF Flow Cytometry Core, supported by the Diabetes Research Center grants NIH P30 DK063720 and NIH S10 1S10OD021822-01.

## Author contributions

Conceptualization, D.R.S., H.W., M.S.A., J.A.B., R.N.G., A.M.; Methodology, D.R.S. and H.W.; Investigation, D.R.S., H.W., J.T.C., Z.L., A.Y., K.P., J.U., A.C.I., J.M.W.; Resources, V.N., J.M.W.; Formal Analysis, D.R.S. and H.W.; Supervision, M.S.A., R.N.G., and A.M.; Funding Acquisition, R.N.G. and A.M.; Writing – Original Draft, D.R.S., H.W., J.T.C., R.N.G., and A.M.; Writing – Review & Editing, D.R.S., H.W., J.T.C., R.N.G., and A.M.

---

**Extended Data Figure 1. Fine mapped T1D association at *IL2RA* locus.** Credible SNPs along with posterior probabilities for primary protective T1D association at the *IL2RA* locus^24^. Expanded sequence view shows the site within the conserved CaRE4 enhancer that harbors the rs61839660 SNP as well as alignment with a MEF2 motif (MEF2_disc1) ^25^ the SNP is predicted to interrupt. The reverse complement of the MEF2 motif is shown for visualization on the top strand of the genomic DNA.

**Extended Data Figure 2. Immunophenotyping of *Il2ra* CaRE3 enhancer deletion on C57BL6 background. a**, Representative CD4 and CD8 staining of thymocyte populations gated on Live CD45+ cells in thymus. **b,** Quantification of normalized percentage of major thymocyte populations, DN (live CD45+CD4−CD8−), DP (live CD45+CD4+CD8+), CD4SP (live CD45+CD4+CD8−), and CD8SP (live CD45+CD4−CD8+). **c**, Histogram of IL2RA surface expression on double negative (CD4−CD8−) thymocytes. **d**, Quantification of IL2RA surface expression on DN thymocytes. **e**, Quantification of abundance of Treg progenitor (FOXP3 TRP: CD4+CD8−FOXP3+IL2RA− and IL2RA TRP: CD4+CD8−IL2RA+FOXP3−) and mature Treg (CD4+CD8−FOXP3+IL2RA+) populations in thymus from CaRE3 EDEL animals **f**, Representative Treg staining in CD4+ T cell compartment from inguinal LN. **g**, Histogram showing IL2RA surface expression on FOXP3+CD4+ Tregs in inguinal LN. **h**, Quantification of Treg percentages of CD4+ T cells in peripheral tissues, including blood, spleen, inguinal LN (iLN), and mesenteric LN (mLN). **i**, Quantification of Il2ra surface expression on Tregs (CD4+FOXP3+IL2RA+) in peripheral tissues. **j**, Quantification of percent IL2RA− Tregs in peripheral tissues. **k**, Quantification of activated Tconv cells (CD4+FOXP3−CD69+) in peripheral tissues. WT (n=5) and CaRE3 EDEL (n=6) littermates were used for experiments. All data are presented as mean +/− s.d. and are representative of at least two independent experiments. *P ≤ 0.05, **P ≤ 0.01, ***P ≤ 0.001, ****P ≤ 0.001 by two-way ANOVA with Sidak’s multiple comparisons test.

**Extended Data Figure 3. Immunophenotyping of *Il2ra* CaRE3 and CaRE4 enhancer deletion mice on non-obese diabetic (NOD) background. a,** Deletion of the conserved *Il2ra* CaRE3 enhancer in NOD mice verified by Sanger sequencing. **b,** CaRE3 deletion Representative thymocyte populations stained for CD4 and CD8 of Live CD45+ cells in CaRE3 EDEL NOD/ShiLt\J mice. **c,** Quantification of percentage of major thymocyte populations, DN (live CD45+CD4−CD8−), DP (live CD45+CD4+CD8+), CD4SP (live CD45+CD4+CD8−), and CD8SP (live CD45+CD4−CD8+). **d,** Representative IL2RA surface expression on DN thymocytes from CaRE3 EDEL NOD mice. **e,** Quantification of percent IL2RA− DN thymocytes and IL2RA levels on IL2RA+ DN thymocytes on CaRE3 EDEL animals. **f,** Quantification of abundance of Treg progenitor (FOXP3 TRP: CD4+CD8-CD73-FOXP3+IL2RA− and IL2RA TRP: CD4+CD8−CD73−IL2RA+FOXP3−) and mature Treg (CD4+CD8−CD73−FOXP3+IL2RA+) populations in thymus from CaRE3 EDEL animals^26^. **g,** Representative FOXP3 and IL2RA staining on CD4+ T cells from inguinal LN of CaRE3 EDEL animals. **h,** Quantification of Treg percentages in peripheral lymphoid organs from CaRE3 EDEL and WT animals. **i,** IL2RA surface expression on Tregs from inguinal LN of CaRE3 EDEL animals. **j,** Quantification of IL2RA levels on IL2RA+FOXP3+ Tregs in CaRE3 EDEL animals. **k,** Quantification of percent IL2RA− Tregs in CaRE3 EDEL animals. **l,** Quantification of activated Tconv (CD4+FOXP3−CD69+) in spleen and inguinal LN. **m**, Time course IL2RA expression in anti-CD3/CD28 activated naïve CD4+ T cells from CaRE3 EDEL and CaRE4 EDEL animals. **n**, Quantification of naïve CD4+ T cell activation showing percent IL2RA− cells. **o,** Quantification of naïve CD4+ T cell activation showing IL2RA MFI on IL2RA+ cells. Data in a-k from CaRE3 (n=3) and WT (n=3) littermate controls. Quantification in m and n are from WT (n=2) and EDEL (n=2) for CaRE3 and CaRE4. Immunophenotyping was performed on male and female mice that were tested to have normal blood glucose levels. All data are presented as mean +/− s.d. and are representative of at least two independent experiments. *P ≤ 0.05, **P ≤ 0.01, ***P ≤ 0.001, ****P ≤ 0.001 by two-way ANOVA with Sidak’s multiple comparisons test.

**Extended Data Figure 4. Islet characterization of NOD CaRE3 and CaRE4 animals. a,** Representative immunofluorescence staining for insulin (green) and CD45 (red) from CaRE4 EDEL (n=1) and WT (n=1) littermate mice. **b,** Representative images showing scoring scale for islet infiltration. **c,** Insulitis quantification in 16-week female WT and CaRE4 EDEL NOD mice. **d,** Insulitis quantification in 32-34-week female WT and CaRE4 EDEL NOD mice. **e,** Insulitis quantification of NOD CaRE4 EDEL mice after anti-PD1 treatment. **f,** Insulitis quantification in 10 week female WT and CaRE3 EDEL NOD mice.

**Extended Data Figure 5. *IL2RA* enhancer deletions reciprocally tune the local competition for IL-2 between self-activated CD4**^+^ **Tconvs and Tregs. a,** *In situ* quantification of pSTAT5^+^ CD4+ Tconvs (CD3+CD4+FOXP3−pSTAT5+) and pSTAT5+ Tregs (CD3+CD4+FOXP3+pSTAT5+) within the panLN paracortex of 6-week-old female mice. panLNs were compared between paired between CaRE4 EDEL (n=3) or CaRE3 EDEL (n=3) versus WT NOD (n=3) littermate controls. Data are from two independent experiments. **b-d,** *In situ* multiparameter analysis of the number of Tregs (x-axis) and CD4+PD1+ Tconvs (y-axis) surrounding individual CD4+PD1+ Tconvs as in Fig. 3. Measurements were made at 30, 40, 50 μm (b through d) away from each individual CD4+PD1+ Tconv.

## Methods

### Mouse Generation

CaRE3 enhancer deletion mice were generated at UCSF by electroporating Cas9 RNPs into mouse blastocysts^27^. Briefly, equal volumes of 160uM CaRE3 1-4 crRNA (IDT) were mixed. An equal volume of 160uM tracrRNA was then added and the gRNA components were allowed to hybridize by incubating at 37C for 10 minutes. We then added an equal volume of 40uM Cas9 protein (QB3 Macrolab) and incubated the solution at 37C for 10 minutes to allow the Cas9 RNP to complex. Mouse lines were established by backcrossing founders to wild type animals at least one generation before performing experiments. Genotyping was done by PCR to look for the expected enhancer deletion. In total four founder lines were established, one on the NOD/ShiLtJ background and three on the C57BL/6J background. All founders were immunophenotyped and phenotypes were found to be consistent. CaRE4 enhancer deletion mice were generated by the Jackson Laboratory and were reported previously^5,8^.

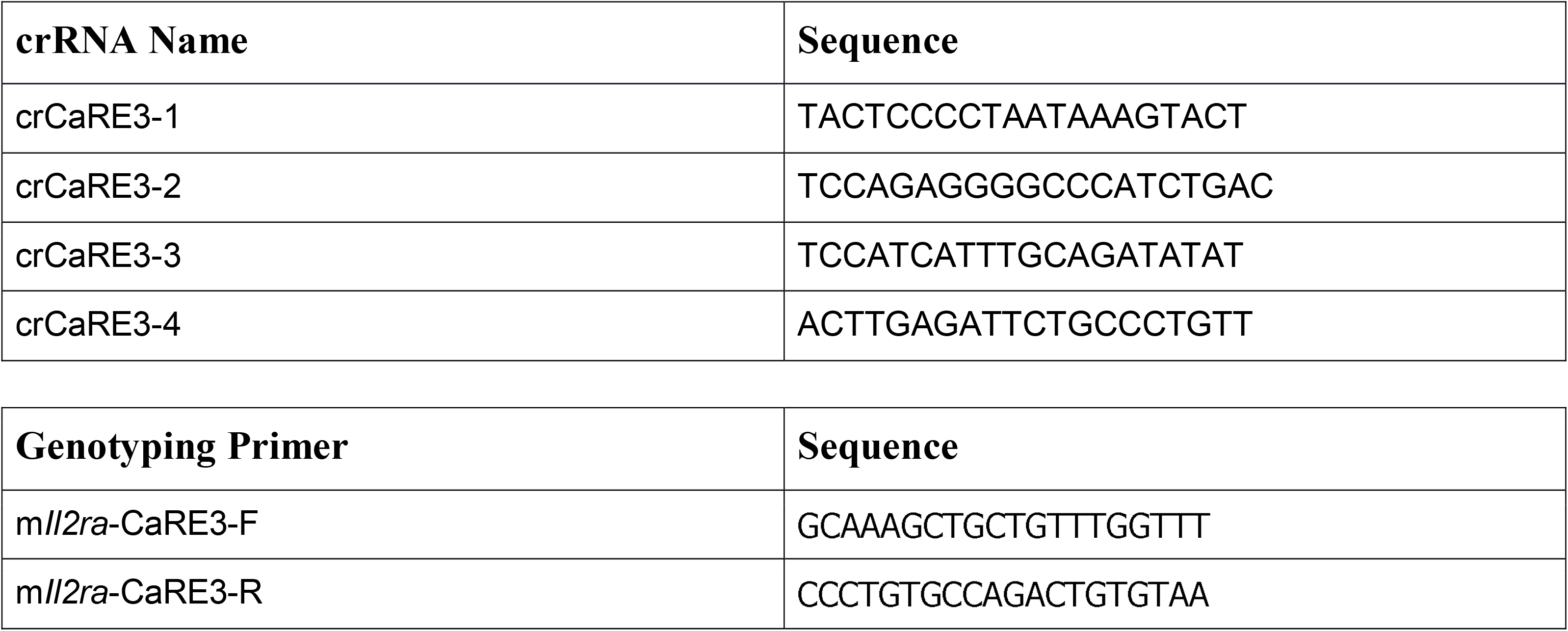

### Mouse experiments and data analysis

All mice were maintained in the UCSF-specific pathogen-free animal facility in accordance with guidelines established by the Institutional Animal Care and Use Committee and Laboratory Animal Resource Center. Experiments were done with animals aged between 1 and 4 months. Wildtype littermate mice were used as controls for all immunophenotyping experiments. All mice used for experiments were normoglycemic unless otherwise noted. Data was excluded only for experimental reasons, either due to failed controls or insufficient numbers for meaningful quantification. Power calculations were not performed and data were assumed to be normally distributed. Experiments were done without blinding or randomization. All data are derived from at least two independent experiments unless otherwise stated. Both male and female NOD mice were used for immunophenotyping experiments.

### Mouse Immunophenotyping

#### Staining

Staining protocol of mouse cells were previously described^5^. In short, staining was done in V-bottom 96 well plates. Surface staining was performed for 10-30 minutes on ice. We followed manufacturer’s protocols for fixation, typically 30 minutes at room temperature. Intracellular staining was performed for 30-60 minutes at room temperature. Antibodies were used at 1:100 dilution. The following staining panels were used in this study:

Thymus

*Surface and Viability:*

Live/Dead-UVB (Invitrogen)

CD45-BUV395 (30-F11)

CD8-PerCP-Cy5.5 (53-6.7)

CD4-BV605 (RM4-5)

CD25-PE-Cy7 (PC61.5)

CD69-APC (H1.2F3)

TCRB-PB (H57-597)*

CD73-PE*

*Intracellular:*

FOXP3-FITC (FJK-16s)

\**not always included*

Spleen and inguinal LN:

*Surface and Viability:*

Live/Dead-UVB (Invitrogen)

CD45-BUV395 (30-F11)

CD8-PerCP-Cy5.5 (53-6.7)

CD4-BV605 (RM4-5)

CD25-PE-Cy7 (PC61.5)

CD69-APC (H1.2F3)

*Intracellular:*

FOXP3-FITC (FJK-16s)

For experiments that required culture of cells complete RPMI with 10% FBS, HEPES, sodium pyruvate, non-essential amino acids, penicillin streptomycin, and beta-mercaptoethanol. For Il2ra time course naïve CD4+ T cells were enriched from splenocytes using EasySep kit (StemCell Technologies). Approximately 100,000 cells were activated using 2ug/ml plate bound anti-CD3/CD28. Cells were stimulated for three days and surface Il2ra expression was measured by flow cytometry every day.

#### Splenocyte Stimulations

Splenocytes were plated at approximately 4 million cells per 500ul of complete RPMI in a well of a 48 well plate. Cells were activated using 1ug/ml anti-CD3 (Clone: 145-2C11, Biolegend) and anti-CD28 (Clone: 37.51, Biolegend). Cells were stained and analyzed by flow cytometry at the end of the time course using the following antibody panels:

Antibodies:

Live/Dead-UVB (Invitrogen, Cat#: L23105)

CD45-BUV395 (30-F11)

TCRB-BV421 (H57-597)

IL2RA-PE-Cy7 (PC61.5)

CD4-BV605 (RM4-5)

CD8-PerCP-Cy5.5 (53-6.7)

FOXP3-FITC (FJK-16s)

### Spontaneous Diabetes

Litters of female NOD mice from CaRE3 and CaRE4 mice were set aside for spontaneous diabetes studies. Blood glucose was measured once a week from a drop of blood at the end of the tail. We measured blood glucose in animals starting at 8-11 weeks of age until 30 weeks of age. Once animals entered the study they were not removed for any reason. The diabetes endpoint was defined as two consecutive measurements of blood glucose over 200mmol/L. Diabetic animals were euthanized according to institutional standards.

### Anti-PD1 Induced Diabetes

For this study we followed a previously published dosing regimen^17^. Briefly, anti-PD1 (Clone: RMP1-14, BioXCell, Cat: BE0146) was administered intraperitoneally to 8-10 week old female NOD mice. Animals were initially dosed with 500ug of antibody (day 0), and then an additional five injections of 250ug each were given every other day (days 2, 4, 6, 8, 10). Mice were followed for a total of 30 days. The diabetes endpoint was defined as two consecutive measurements of blood glucose over 200mmol/L. Diabetic animals were euthanized according to institutional standards.

### Pancreas Histology for Immune Infiltration

Briefly, the pancreas was isolated and placed into zinc formaldehyde fixative overnight at 4C. The next day the tissue was washed with water, 5 minutes per wash with gentle rocking. Three more washes were done with 1x PBS in a similar manner. Tissue was transferred to 70% EtOH and stored at 4C before further processing. Pancreas processing from 32-week-old animals was done by the UCSF Histology Core (Diabetes Center, UCSF). H&E stained sections were mounted on slides and scored under a microscope. Pancreas processing for 16-week-old animals was done by HistoWiz which completed embedding, sectioning, H&E staining, and imaging. Pancreatic islets were categorized as being normal, or as peri-insulitis or insulitis depending on severity and overall appearance of immune infiltration. Examples of this are shown in Extended Data Fig. 4.

### Tissue section preparation, processing and immunostaining

pLNs were isolated from 6-week old female NOD mice and fixed for 14h at 4°C in BD Cytoperm/Cytofix (BD Bioscience, Cat#: 554722) diluted 1/4 in PBS. pLNs were washed 3x in PBS (10 min per wash), carefully trimmed of fat using a stereo dissection microscope and fine forceps, and dehydrated for 24 h in a 30% sucrose solution made in 0.1 M phosphate buffer. LNs were then embedded in optimal cutting temperature (O.C.T.) compound (Sakura Finetek, Cat#: 50-363-579), frozen on dry ice, and stored at −80°C. 18-50 μm sagittal LN sections were prepared using a cryostat (Leica) equipped with a 40 Surgipath DB80LX blade (Leica, Cat#: 14035843497). Cryochamber and Specimen cooling was set to −17°C.

Tissue sections were adhered to Superfrost Plus microscopy slides (VWR, Cat#: 48311-703), permeabilized using 0.1% Triton X-100 for 10 min at 22°C, blocked in 5% mouse serum for 1 h at 22°C, and washed in PBS. Tissue sections were next incubated with directly conjugated primary antibodies diluted in PBS for 15 h at 4°C. After washing 3x in PBS (10 min per wash) at 22°C, samples were mounted in Fluoromount-G (SouthernBiotech, Cat#: 0100-01), which was allowed to cure for a minimum of 14h at 22°C. All imaging was performed using No. 1.5 coverglass (VWR, Cat#: 48393-241). The following directly-conjugated antibodies were used for immunostaining: anti-CD3 (clone 17A2, 1 μg/ml), anti-CD4 (clone RM4-5, 1 μg/ml), anti-FOXP3 (clone FJK-16s, 1 μg/ml), anti-IL2RA (clone PC61, 1 μg/ml), anti-PD1 (clone 29F.1A12, 1 μg/ml). Combinations of the following organic fluorophores were used: Brilliant Violet 421, Brilliant Violet 480, Alexa Fluor 555, eFluor 570, Alexa Fluor 594, eFluor660, Alexa Fluor 647, Alexa Fluor 488, and Alexa Fluor 700.

For pSTAT5 immunostaining, fixed tissue sections were permeabilized in pre-chilled 100% methanol for 18 min at −20°C, washed extensively in PBS, blocked in 5% donkey serum for 1 h at 22°C, and washed further in PBS. Tissue sections were next incubated with unconjugated anti-pSTAT5 (clone C11C5, 1.715 μg/ml) diluted in PBS for 15 h at 4°C. Following washing in PBS at 22°C, tissue sections were incubated with either Alexa 488, Alexa 594, or Alexa 647-conjugated donkey anti-rabbit F(ab’)2 fragments (0.483 μg/ml, Jackson ImmunoResearch Laboratories, Inc, Cat#: 711-546-152 or 711-6060152) for 2h at 22°C. Sections were then washed 4x in PBS (10 min per wash) at 22°C, stained with directly-conjugated antibodies as described above, and mounted in Fluoromount-G.

### Laser scanning confocal microscopy

Digital images were acquired using an upright Leica TCS SP8 X spectral detection system (Leica) equipped with a pulsed white light laser, 4 Gallium-Arsenide Phosphide (GaAsP) Hybrid Detectors (HyDs), 1 photomultiplier tube (PMT), 40x (NA = 1.3) and a motorized stage. For tissue sections (18-20 μm), images were acquired with a pixel size of 0.271-0.286 μm, z step size of 0.3-1 μm, and detector bit-depth of 12.

### Image processing and segmentation

Image files generated in LAS X software were converted into “.ims” files in Imaris software (Bitplane) and subjected to a 1 pixel Gaussian filter to reduce noise.

Segmentation of densely packed T cells was performed using a protocol modified from Li et al. and described in Wong et al.^19,28^. In brief, this process involved creating artificial T cell nuclei, which in turn were subjected to segmentation algorithms. Initially, “.ims” files were imported into Fiji^29^. The brightness and contrast of the CD4 channel was adjusted linearly to thinly demarcate the plasma membranes of T cells. The adjusted CD4 channel was then converted into an 8-bit format, duplicated and inverted to create an *inverted sum* channel. Next, the original channel was subtracted from the *inverted sum* channel, producing a *high-contrast inverted* channel, which was subsequently binarized using the “Auto Local Thresholding” tool. Binarized images were then despeckled to remove noise and subtracted from the *inverted sum* channel to improve separation between artificial T cell nuclei. The final product was exported as a “.TIFF” image sequence, imported into the original “.ims” file in Imaris, and subjected to the “Surface Object Creation” module. Segmentation artefacts were excluded using a combination of sphericity and volume thresholds, as well as manual correction.

### Statistics

Statistical calculations were performed using GraphPad prism versions 7.0 and 8.0. Tests between two groups used a two-tailed Student’s t-test. Tests between multiple groups comparing EDEL to WT littermate controls used two-way ANOVA with Sidak’s test for multiple comparisons.

## Code Availability

The code used for spatial image analysis of immunostained panLNs in this study has been deposited on GitHub (https://github.com/harikesh22/Treg_neighbourhood_analysis). There are no restrictions on code availability.

## Data availability

All data generated or analyzed are included in the published Article and the Supplementary Information. GWAS SNP tracks in Figure 1a derived from papers referenced in figure legend. H3K27Ac ChIP-seq data in Figure 1b for Tnaive (E039 Primary T helper naïve cells from peripheral blood), Tstim (E041 Primary T helper cells PMA-I stimulated), and Treg (E044 Primary T regulatory cells from peripheral blood) was generated as part of the Epigenome Roadmap Project (https://egg2.wustl.edu/roadmap/web_portal/). The FOXP3 and STAT5 ChIP-seq data in Figure 1b is publicly available as part of the FANTOM5 project. Data was uploaded to UCSC Browser for visualization and export (http://www.ag-rehli.de/TrackHubs/hub_Tsub.txt). There are no restrictions on data availability.

