## Supplementary figures and images for "T cell subset-selective *IL2RA* enhancers shape autoimmune diabetes risk"

### Extended Data 1

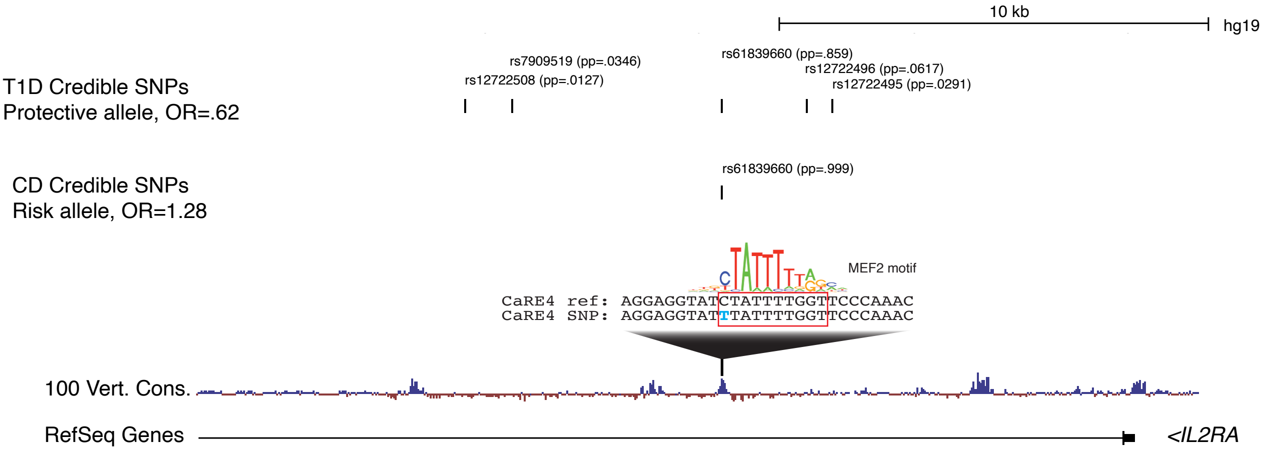

### Extended Data 2

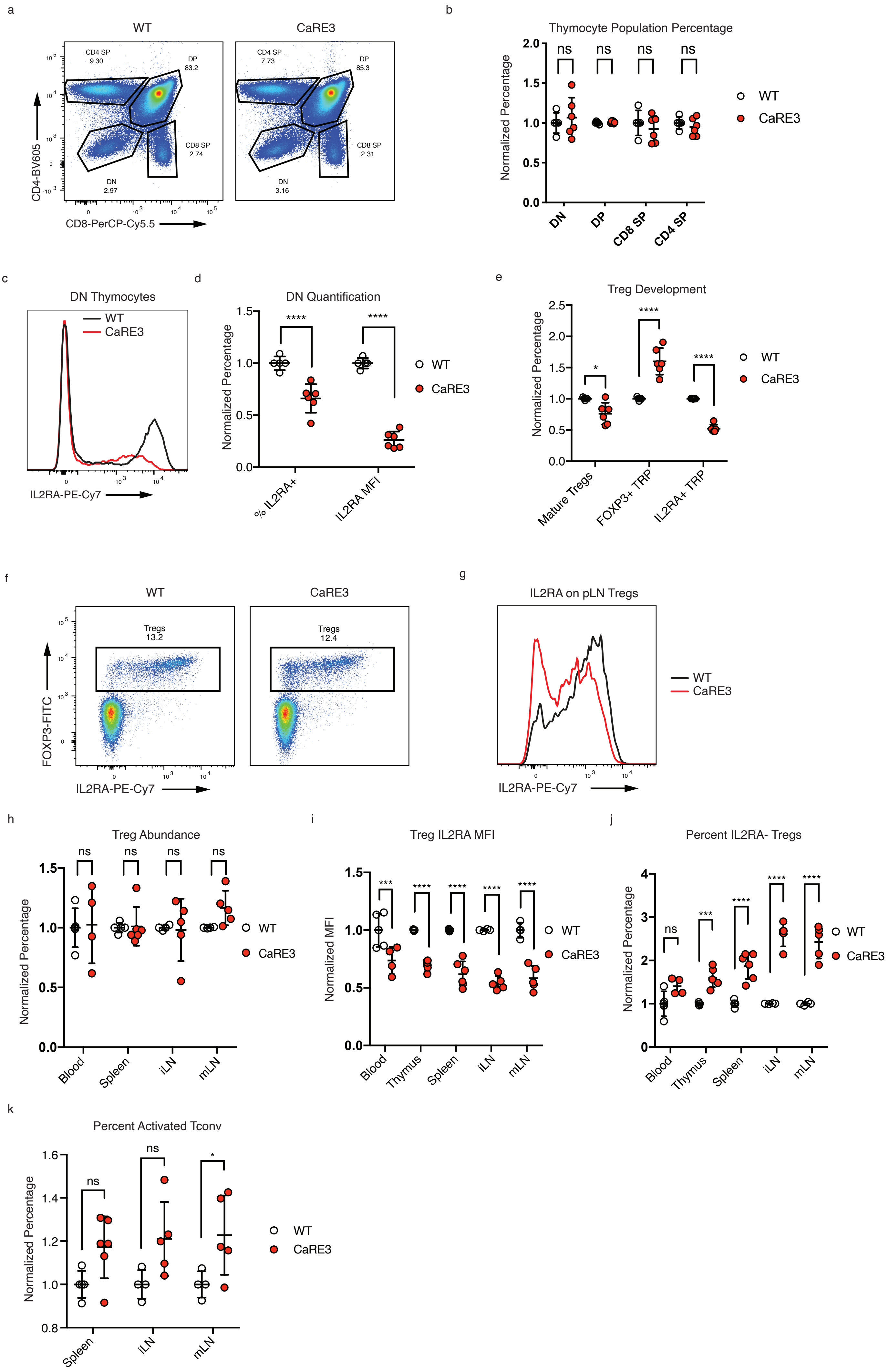

### Extended Data 3

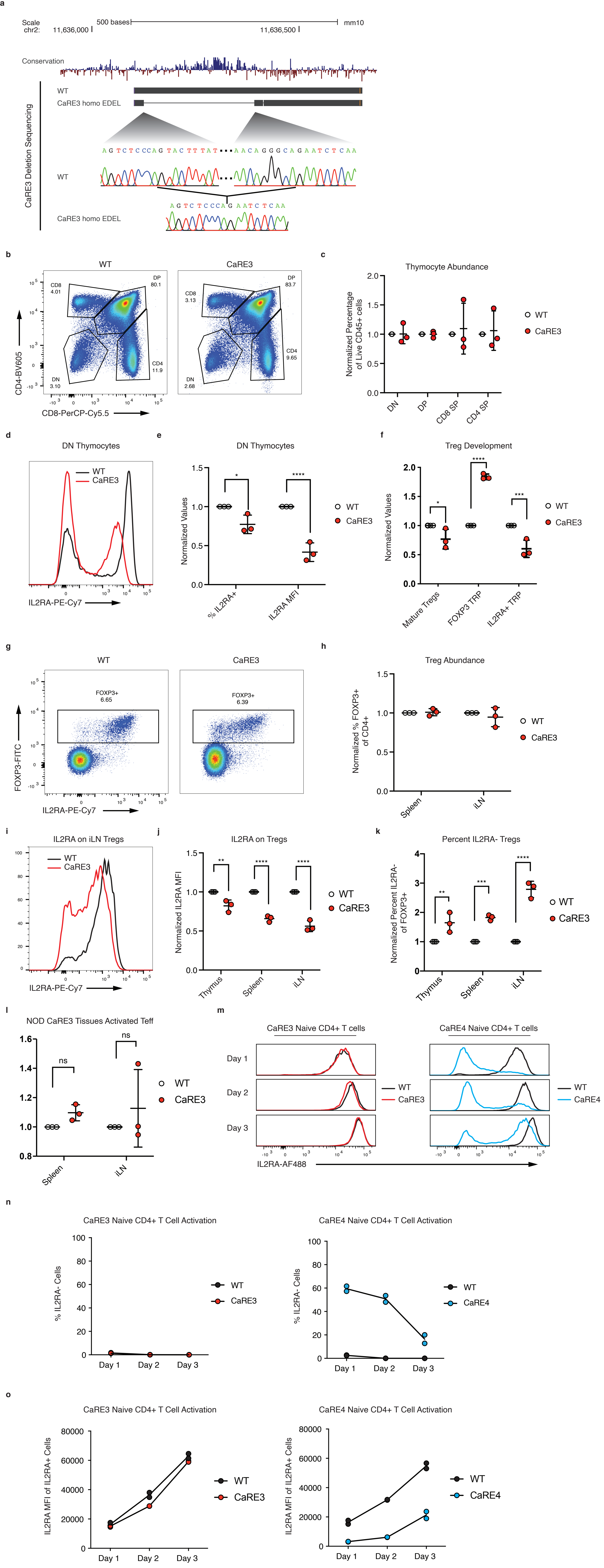

### Extended Data 4

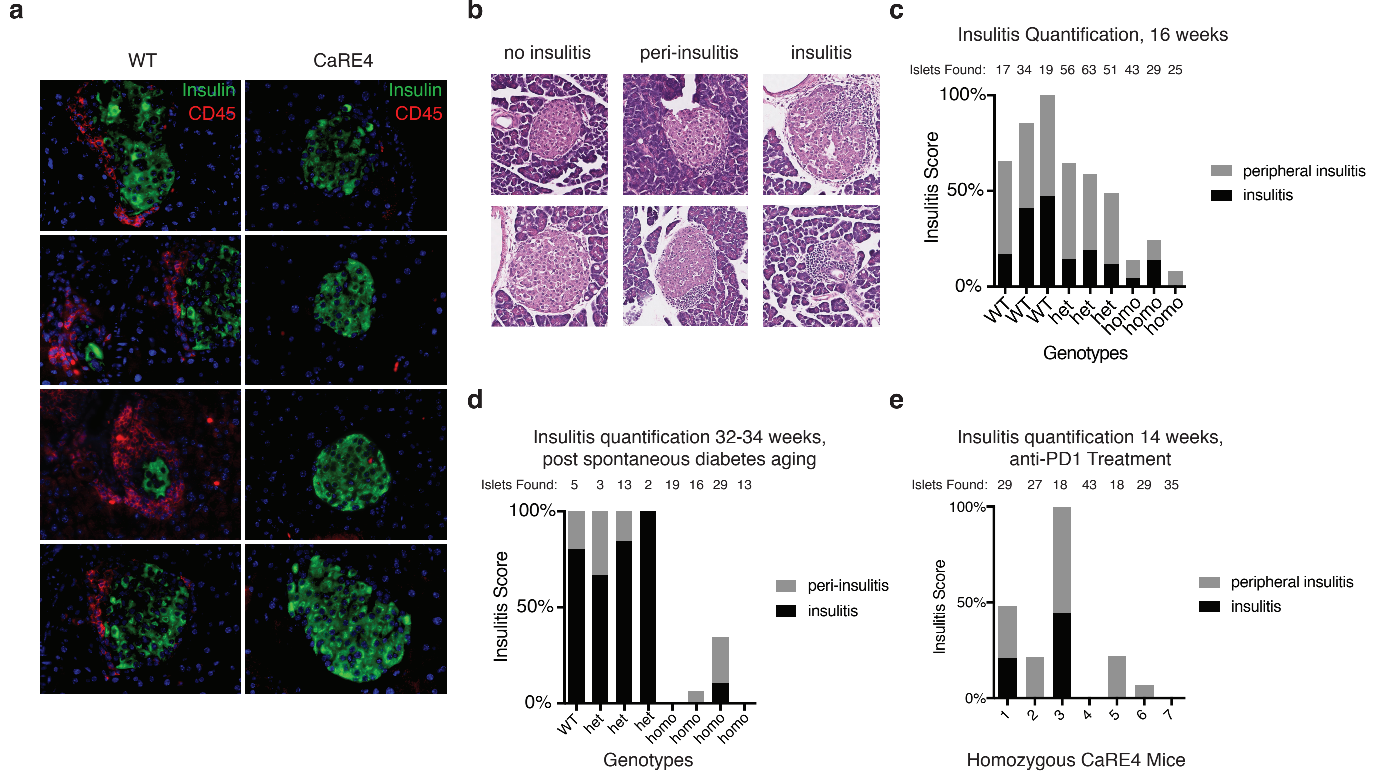

### Extended Data 5

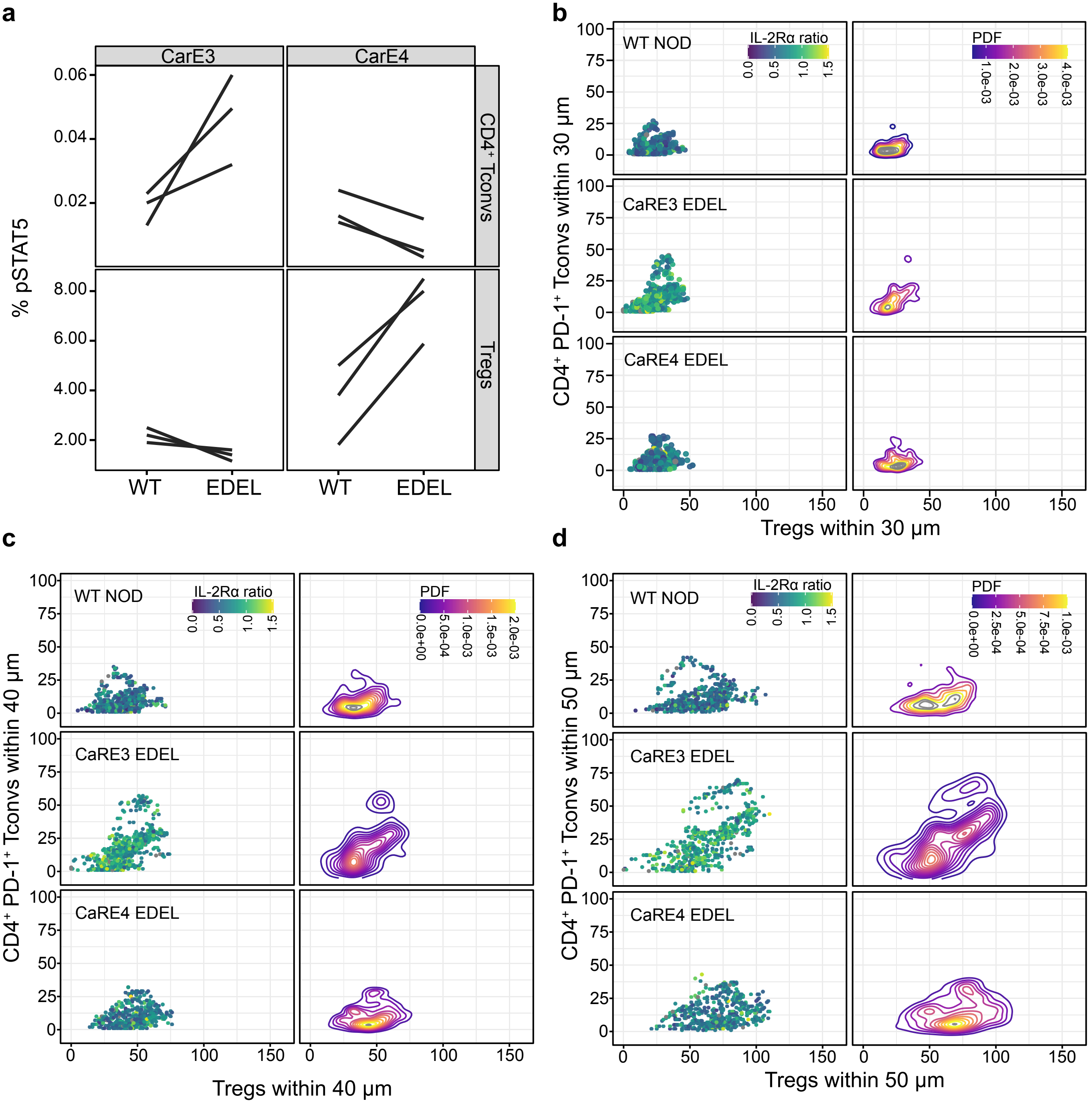
